# Estimation of immune cell content in tumor using single-cell RNA-seq reference data

**DOI:** 10.1101/663500

**Authors:** Xiaoqing Yu, Y. Ann Chen, Jose R. Conejo-Garcia, Christine H. Chung, Xuefeng Wang

## Abstract

**Background:** The rapid development of single-cell RNA sequencing (scRNA-seq) provides unprecedented opportunities to study the tumor ecosystem that involves a heterogeneous mixture of cell types. However, the majority of previous and current studies related to translational and molecular oncology have only focused on the bulk tumor and there is a wealth of gene expression data accumulated with matched clinical outcomes.

**Results:** In this paper, we introduce a scheme for characterizing cell compositions from bulk tumor gene expression by integrating signatures learned from scRNA-seq data. We derived the reference expression matrix to each cell type based on cell subpopulations identified in head and neck cancer dataset. Our results suggest that scRNA-Req-derived reference matrix outperforms the existing gene panel and reference matrix with respect to distinguishing immune cell subtypes.

**Conclusions:** Findings and resources created from this study enable future and secondary analysis of tumor RNA mixtures in head and neck cancer for a more accurate cellular deconvolution, and can facilitate the profiling of the immune infiltration in other solid tumors due to the expression homogeneity observed in immune cells.

## BACKGROUND

Cancer immunotherapy has made substantial progress and has dramatically impacted the treatment of multiple cancers, including skin cancer, lung cancer, and head and neck cancer. The cellular composition of a tumor and its immune microenvironment varies between patients and tissue types. The presence and higher content of tumor-infiltrating lymphocytes (TILs) is believed to be associated with response to the immunotherapy. In melanoma, it was also found that the composition of immune cells such as CD8+ cytotoxic lymphocytes and dendritic cells are strong prognostic predictors themselves and are associated with overall clinical outcomes. However, there are still considerable technological and analytical barriers to assess cancer and immune cell compositions in the tumor quantitatively. The pathological approaches such as immunohistochemical (IHC) staining and flow cytometry analysis are labor intensive and often involve considerable inter-observer variation. Therefore, the cell decomposition based on existing molecular profiles of tumors has received many attentions in recent years. Earlier work has been centered on whole exome sequencing data. Based on DNA mutational signatures and the distribution of local copy numbers, several methods have been proposed to infer the tumor purity—defined as the proportion of cancerous cells in the tumor tissue. Based on the similar computational model (Carter 2012), subclonal heterogeneity and somatic homozygosity can also be explored. Previous studies have also attempted to deconvolve gene expression profiles (including microarray and RNA-seq) of tumor samples to infer the stromal and immune cell admixture (Yoshihara 2013). These methods leverage distinct transcriptional properties of different cell types, which provide finer granularity in the cell composition estimation than using DNA mutational profiles alone.

The software CIBERSORT has now been widely used in the area to estimate immune cell subsets from tumor expression profiles. But its application has been limited to microarray studies due to the source of the training gene expression panel. Only recently have efforts begun to extend the cell deconvolution method to RNA-seq data and to identify more microenvironment-informative markers. These reference markers were selected from whole transcriptome data and narrowed down through correlating gene expression with tumor purity estimates. The nCounter system (NanoString) has gained popularity in the clinical and translational setting as an alternative tool for immune cell profiling. The advantage of NanoString platform is that it is based on a highly sensitive and non-enzymatic process to enable a more precise quantification of RNA expression, which provides reliable data even with FFPE samples. However, nCounter is a targeted gene expression panel, and the surrogate expression profile cannot differentiate all cell subpopulations. Therefore, there is a pressing need to develop more efficient gene reference panel and related computational tools to quantify the components of tumor microenvironment in situ on a larger scale, which will facilitate both retrospective and prospective studies.

The recent maturation of single-cell RNA sequencing (scRNA-seq) has enabled us to directly profile the cell composition and understand tumor heterogeneity at a cellular level. With newly developed high-throughput cell sorting and barcoding technologies, thousands of individual cells per tumor can be profiled in parallel to capture intra-tumor heterogeneity at an unprecedented resolution [1–3]. Unless the main goal of a project is to study underrepresented cell populations, scRNA-seq experiments can be done without the need for cell sorting--which is laborious and prone to considerable bias due to cell death and cell selection. The unbiased and simultaneous characterization of both immune and cancer cell is essential for tracking and forecasting the tumor ecosystem, e.g., in patients before and after immunotherapy. The cellular composition, as well as the relationships between different cell subpopulations, are generally explored by clustering analysis using all gene expression data--most notably, based on the method called t-Distributed Stochastic Neighbor Embedding (t-SNE). Cell types corresponding to each cell cluster can then be inferred based on existing cell-type-specific marker genes and any available prior knowledge about the cells. Furthermore, a differential expression analysis between distinct cell populations may provide new marker genes for cell mixture deconvolution. Nevertheless, large-scale scRNA-seq studies involve expensive sequencing efforts, prohibiting them from being more widely used in practical and clinical settings. There is still considerable interest in the community to drive cell-type-informative markers for facilitating the analysis of bulk tumor sequencing. It thus motivates us to derive more efficient cell-type-informative markers by leveraging high-quality scRNA-seq data generated from existing studies.

Here we investigated gene expression profiles of 6,000 single cells from 15 head and neck squamous cell carcinoma (HNSCC) patients. To allow for a finer deconvolution of immune cell subtypes, we employ an adaptive divide-and-conquer scheme to isolate cell populations in silico. The reference gene expression profile matrix was then built based on identified single cell populations. We show that the reference profiles obtained from single cell expression data enable a more reliable estimation of cellular composition in bulk tumor, and they have ability to discriminate immune cell types with finer granularity. Our work demonstrates that established single cell gene expression in each tumor type can further add value to the digital dissection the tumor microenvironments. We provide these reference matrices and gene panels, namely single-cell gene expression profiles (scGEPs), to the community as a useful resource for studying heterogeneous tumor ecosystems.

## METHODS

### Single-cell RNAseq Data

We downloaded the single-cell RNA-seq data from Puram et al. [1] which generated expression data of 6,000 single cells from head and neck squamous cell carcinoma (HNSCC) patients. By reviewing all published single-cell RNA-seq data (up to Dec 2018) in cancer, we found that this dataset covered the most diverse stromal, malignment and immune cells in the tumor microenvironment (TME), and relatively large number of patients. Importantly, it provides annotated cells from four T cell major subpopulations: regulatory T-cells (Tregs), convectional CD4+ T cells (CD4^+^ Tconv), CD8^+^ T and CD8^+^ T exhausted. Therefore, single cell expression profiles from Puram et al. study is an ideal source of reference data. Note that expression profiles of malignant cells are highly specific to HNSCC, but we hypothesize that expression reference of immune cells is applicable to other cancer types. After removing the patient samples with less than 50 cells, 5712 cells from 16 treatment-naive patients plus matched lymph nodes from three of these patients remained for analysis (Table S1). As described in Methods in Puram et al. [1] gene expressions were quantified as y=log2 (TPM^+^1), where TPM refers to transcripts per million, a gene quantification method that has been considered superior to FPKM (fragments per kilobase per million read) and more robust to differences in RNA library size [4].

### Enrichment analysis of cell-type-specific genes

We adapted the single-sample Gene Set Enrichment Analysis, or ssGSEA [5], to calculate the enrichment scores of pre-existing cell-type-specific marker genes. These scores will be used to assist the cell type assignment step to be described in the following sections. ssGSEA is an extension of GSEA method that computes an aggregated enrichment score for a gene set. But instead of gene-phenotype association score, ssGSEA considers rankings of gene expression relative to remaining genes in the genome within each sample, and calculate a score that represents the degree that genes in a gene set are coordinately up-or down-regulated. Signature genes for HNSCC tumor, immune, and stromal cells were obtained from previous studies [1, 6, 7]. To choose the most reliable and generalized signatures, we used only the genes shared by all resources. Together, we collected 140 signature genes covering 15 cell types including HNSCC tumor cells, immune cells, T cell subtypes, and stromal cells. The curated gene list is given in Supplementary Table S2. Note that this list alone is not sufficient to be used as a reference panel for the cell content deconvolution with bulk tumor gene expression data. Enrichment of each cell-type signature was assessed using ssGSEA implemented in R package gsva [6].

### Cell type identification

Similar to the data analysis presented in Purma study, we choose to use the t-SNE method to visualize the cell clusters and explore the cell type compositions based on transcriptomes of all examined cells. However, as shown in the previous analysis and in the results section, t-SNE method alone is only able to identify clusters of major cell types and not able to distinguish between T cell subpopulations. Furthermore, the location of the clusters in the t-SNE map and their relative positions to other clusters will change across analysis runs. As a limitation of the technique, t-SNE cannot reproduce the same clustering map if different cells or perplexity parameters are chosen in one analysis run. Therefore, we propose to use a multi-stage cell identification scheme for obtaining more accurate cell type inference--by adaptively integrating t-SNE and ssGSEA results. The steps and detailed parameters used are described below.

1. Tumor cell classification: To classify HNSCC malignant cells, we performed t-SNE analysis of all cells using perplexity parameter of 50 followed by DBscan clustering (with parameters eps=5 and minPts =5). Clusters were classified as malignant cells and non-malignant cells based on their ssGSEA enrichment scores using signature genes for HNSCC tumor cells (Figure S1A,B). As reported previously in various cancer studies [1, 3], malignant cells were clustered by patients while non-malignant cells were clustered by cell types (Figure S1C).
2. Non-tumor cell classification: The non-tumor cells identified in step 1 were subjective to a secondary stage of clustering analysis. t-SNE with the perplexity of 30 was performed followed by DBscan clustering (with parameters eps=6 and minPts =15). These parameters were chosen based on two criteria: (1) the resulted clusters should maximize the degree of differentiation of cell populations; (2) the resulted clusters should have the greatest consensus possible with the ssGSEA metrics. Based on the ssGSEA enrichment scores, clusters are assigned to major immune and stromal cell types including Fibroblasts, B cell, Macrophages, Endothelial cells, Dendritic cells, Mast cells and T cells (Figure S2A and Figure S2B).
3. T cell subtype identification: Similar procedure was used to classify T cell subtypes from the lumped T cells population identified in step 2. We performed single-cell consensus clustering (SC3) analysis [8] and were able to identify four distinct clusters of T cell subpopulations. These four clusters were assigned to conventional CD4^+^ T cells (CD4^+^ Tconv), T-regulatory cells (Treg), conventional CD8^+^ T cells (CD8^+^ Tconv), and exhausted CD8^+^ T cells, based on their ssGSEA enrichment scores (Figure S3A and Figure S3B). Next, differential expression analysis was performed comparing CD4 Tconv vs. Treg cells, and CD8^+^ Tconv vs. exhausted CD8^+^ T cells using R package limma [9]. Only genes with |log2FoldChange| >1 and Benjamini-Hochberg adjusted p-value < 0.05 were considered significantly differentially expressed and reported in Table S3. The identified differentially expressed genes were compared with previously reported marker genes for these cell types.

### scRNA-derived marker genes

To develop a finer panel of cell-type-specific genes, we identified marker genes that are specifically expressed in each cell type. Differential expression analysis was first performed between any pairs of the 11 cell types using R package limma. Then marker genes of each cell type were identified as those significantly highly expressed in cell type under consideration compared to at least 5 other cell types (log2FoldChange >3 and Benjamini-Hochberg adjusted p-value < 0.05). In total, we identified 581 marker genes and reported the gene names and limma results in Table S4.

### Deconvolution method for bulk tumor

The objective of the deconvolution algorithm is designed to solve for the linear equations **m** = **f** x **B**, where m is the input gene expression profile (GEP) matrix, **f** is a vector of cell fractions to be estimated, and **B** is the gene expression signature or reference GEP matrix. A machine learning method, v-support vector regression (v-SVR) combining feature selection with a linear loss function and L2-regularisation [10], was used to infer the compositions of the malignant cells, tumor-infiltrating cell types/subtypes, and stromal cells from the bulk gene expression. This method has been implemented in CIBERSOR [11], a tool that has now been widely used for in cancer research. The initial setting of CIBERSORT was designed for estimating 22 immune cell types using 547 signature genes (LM22) derived from microarray data. In this study, we will apply the same SVR method implemented in CIBERSORT to infer cell types that are more representative in head and neck tumors. The reference GEP panels used in SVR will be described in the following section.

### In silico assessment of final reference GEP panels

With the availability of high-resolution scRNA-seq data, one main objective of this study is to explore new ways to generate the reference GEP matrices to be used in bulk tumor deconvolution, i.e., the matrix B as described in the previous section. The ideal B matrix should be able to yield maximal and robust discriminatory power between cell type clusters. Meanwhile, the pooled scRNA-seq data can be served as ground truth for benchmarking the performance of reference GEP as well as deconvolution methods—because the true cell composition in the bulk gene expression data will be known. The similar idea has been implemented in a recent study [7]. The first step of constructing reference GEP matrices is to choose a panel of reference genes that can distinguish the cell populations. In this study, we will focus on four gene panels: (1) LM22 gene reference panel, designed by Newman et al: it contains 547 genes that distinguish 22 human hematopoietic cell phenotypes including several T-cells types, B cells, and natural killer cells. This panel is the default panel used in CIBERSORT and thus has been used extensively; (2) A panel of signature genes identified from previous literature: it contains 140 genes that are served as signatures for 15 major cell types including HNSCC tumor cells, immune cells, T cell subtypes, and stromal cells (Table S2). (3) The scRNA-derived marker gene panel discovered through the steps described previously in the method: which contains genes that uniquely expressed in each cell population identified from HNSC scRNA-seq data (Table S4); (4) A T-cell-specific GEP panel discovered through steps similar to GEP panel (3) but with a focus on four T cell subtypes (Table S3). Note that we only used the gene list information of these panels. The GEP matrix of these genes is formed through averaging all single cells assigned to these populations. In order to assess the prediction performance of the above four GEP panels, we tested them on in silico bulk tumors by aggregating the single cell transcriptome data. Expression data of individual cells from the same patient in Puram study were pooled to form 15 in-silico tumors, which exhibit varied cellular compositions.

## RESULTS

### Identifiable cell types using HNSCC single cell data

Overall, the adaptive clustering analysis on single-cell transcriptome data pooled from all HNSCC tumor samples identified distinct 11 cell clusters to be used in generating reference GEP. These cells types are: HNSCC Malignant cells, Fibroblasts, Macrophages, Dendritic cells, Endothelial cells, Mast cells, B cells, conventional CD4^+^ T cells, T-regulatory cells, conventional CD8^+^ T cells, and exhausted CD8^+^ T cells. As shown in the t-SNE plot with all cells projected (Figure 1A), most cells from same immune cell types are grouped together while malignant cell and Fibroblasts cell clusters contains multiple subgroups within each cluster. In the follow-up analyses, we will show that these subgroups are mainly driven by inter-tumor heterogeneity. The cell grouping information was then used to construct the cell composition map back in each tumor. As illustrated in the stacked bar chart in Figure 1B, the proportions of malignant cells (tumor purity) vary uniformly between 0 and 1. This pattern reflects the original experimental design and is consistent with results from the original analysis [1]. We also observed that some important immune subsets such as tumor-infiltrating Treg cells (coded with dark blue) only exist in tumor samples with lower tumor purity, i.e. sample towards the right side of the plot. Treg cells plays important role as regulators of anti-tumor immune suppression and Treg/CD8^+^ T cell ratio may have a clinical significance in analyzing tumors in HNSCC patients [12]. However, results from scRNA-seq data suggests that the overall Treg expression signature may be underrepresented in genomic projects that are biased towards tumors with higher purity, such as TCGA. In the following, we briefly describe results generated from each step. First, we observed that the unsupervised clustering on all cells based on t-SNE revealed eight major clusters as depicted in Figure S1A. Note that, at this stage, we had no information about cell types underlying these cell groups and the number of clusters might differ subject to the perplexity parameter choice in t-SNE. We started the cell type identification from first distinguishing tumor and non-tumor cells. By adding ssGSEA scores representing the tumor cell signature into the t-SNE map (Figure S1B), we identified two major cluster regions of malignant cells located in the very top and lower regions. By further adding the color layers reflecting the tumor origin, we observed that the cell clusters in these regions were clearly separated by patient IDs while they were mixed together in a mosaic pattern in other cluster regions (Figure S1C). The results above align with previous findings [1, 3, 7] that inter-tumor heterogeneity may arise more at the tumor malignant cell level than at the immune cell level—suggesting that immune cell signatures abstracted from the proposed scheme will be applicable to not only HNSCC samples generated from different studies but also samples from different tumor types. Next, we performed a second round of t-SNE analysis by excluding all tumor cells identified from previous steps. The new clustering analysis revealed seven major cell clusters (Figure S2A). We were able to identify the cell types corresponding to each cluster by adding ssGSEA score specific to Fibroblasts, B cell, Macrophages, endothelial cells, dendritic cells, mast cell, and T cells one at each time as depicted in Figure S2B. As expected, this subset of cell population is dominated by Fibroblasts and T cells. When we adding the color layers reflecting patient origins into Figure S2A, we found a similar pattern that patient IDs were mixed together in each cell type cluster, indicating that the sub-clusters (such as in the T cells) may reveal further cell subtypes. This leads us to the next step by further zooming into the expression profiles of cells from T cell populations.

**Figure 1.**
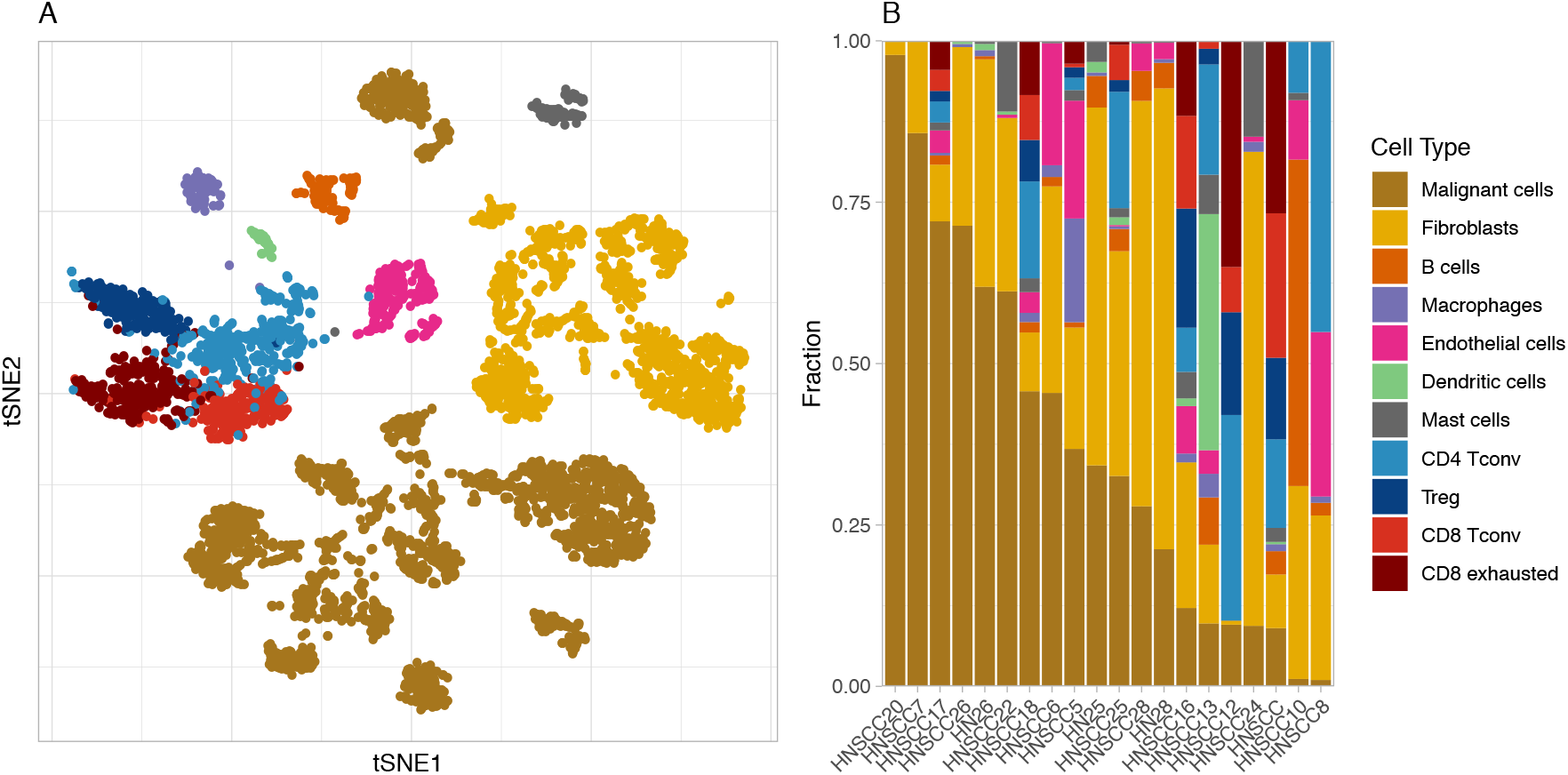
Profiling cellular composition of HNSCC tumors using scRNA-seq. (A) 2D t-sne projection of the expression profiles of 5,712 single cells (3,259 immune cells and 2,453 malignant cells) from 20 HNSCC tumors and lymph node samples of 16 patients. Single cells are shown in dot and colored by cell types. (B) Cell composition per sample. Patients are ordered by their fractions of malignant cells.

### Deconvolution of T cell subtypes using identified T cell population

Based on SC3, we further identified four clusters from T cells (Figures 2A). The cell types in the T cell subpopulation were first determined based on the gene enrichment signatures of CD4^+^ and CD8^+^ cells (as shown in the upper panel in Figure S3B). Within these two subpopulations, [CD8^+^ cells further marked with ssGSEA signatures for CD8^+^ Tconv and CD8^+^ exhausted; and CD4^+^ cells were marked with CD4^+^ Tconv and Treg cells signature values (Figure S3B). As shown in Figure S3, the signatures for two CD8^+^ cell types are overlapped and it is difficult to assign these cells to any subtypes. As further summarized in the heatmap of ssGSEA scores (Figure S4), the ssGSEA analysis based on curated signature genes were able to distinguish between major cell types using single cell level expression data but failed to provide the necessary granularity in separating T cell subtypes. To determine T cell subtypes, especially CD8^+^ subtypes, we performed differential expression analysis between the two cell groups identified within CD4^+^ T cells and CD8^+^ T cells. Differentially expressed genes (adjusted p value < 0.05, limma moderated ř-test, and |log2fold-change| >1) are reported in Table S3. Cell subtypes were then inferred from the status of top differentially expressed genes, by comparing them with existing cell-type-specific marker genes. Figure 2B and Figure 2C are heatmaps depicting top differentially expressed genes between CD8^+^ cell clusters and CD4^+^ cell clusters, respectively. Candidate genes that overlapped with marker genes identified from previous studies are listed and labeled in heatmaps. Note that several exhaustion-related genes can serve as markers for separating both subtypes in CD4 and CD8, such as *TIGIT* and *CTLA4.* For the CD8^+^ T cell subtypes, we compared the candidate marker genes identified in our DE analysis to the exhausted CD8^+^ T cells marker genes reported in a previous single-cell RNA-seq from infiltrating T cells of lung cancer [13]. A total of 36 genes are found shared by the two studies and all labeled in Figure 2B. Among these 36 genes also includes 14 known exhaustion markers, such as *PCCD1, TIGIT, HAVCR2,* and *CTLA4* (Figure 2B, text in red), which further confirmed the identify of these exhausted CD8^+^ T cells. The other CD8^+^ T cell cluster without expression of exhaustion genes is considered as conventional CD8^+^ T cells. For the CD4^+^ T cell subtypes, we also compared the candidate marker genes identified from the DE analysis with the Tregs marker genes reported by four previously published scRNA-seq data from different cancer types [13–16] (Figure 2D). We observed that there were 20 genes shared by all five studies (Figure 2C, text in red), including known Tregs markers *FOXP3, TIGIT,* and *CLTA4;* and there were many more genes previously identified at least once (Figure 2E). Our study also identified 207 genes that uniquely enriched in this HNSCC dataset (Figure 2E), including *PPP1CA, RUNX3, CCR6,* and *PSMB8* which were previously reported to be associated with Tregs and their functions [17–20]. Based on these observations, we assigned Tregs to this cluster of CD4^+^ T cells. The other CD4^+^ cluster with low expression of exhaustion markers and with exclusively high expression of CCR7, CXCR4, and TOBI was considered as conventional CD4^+^ T cells.

**Figure 2.**
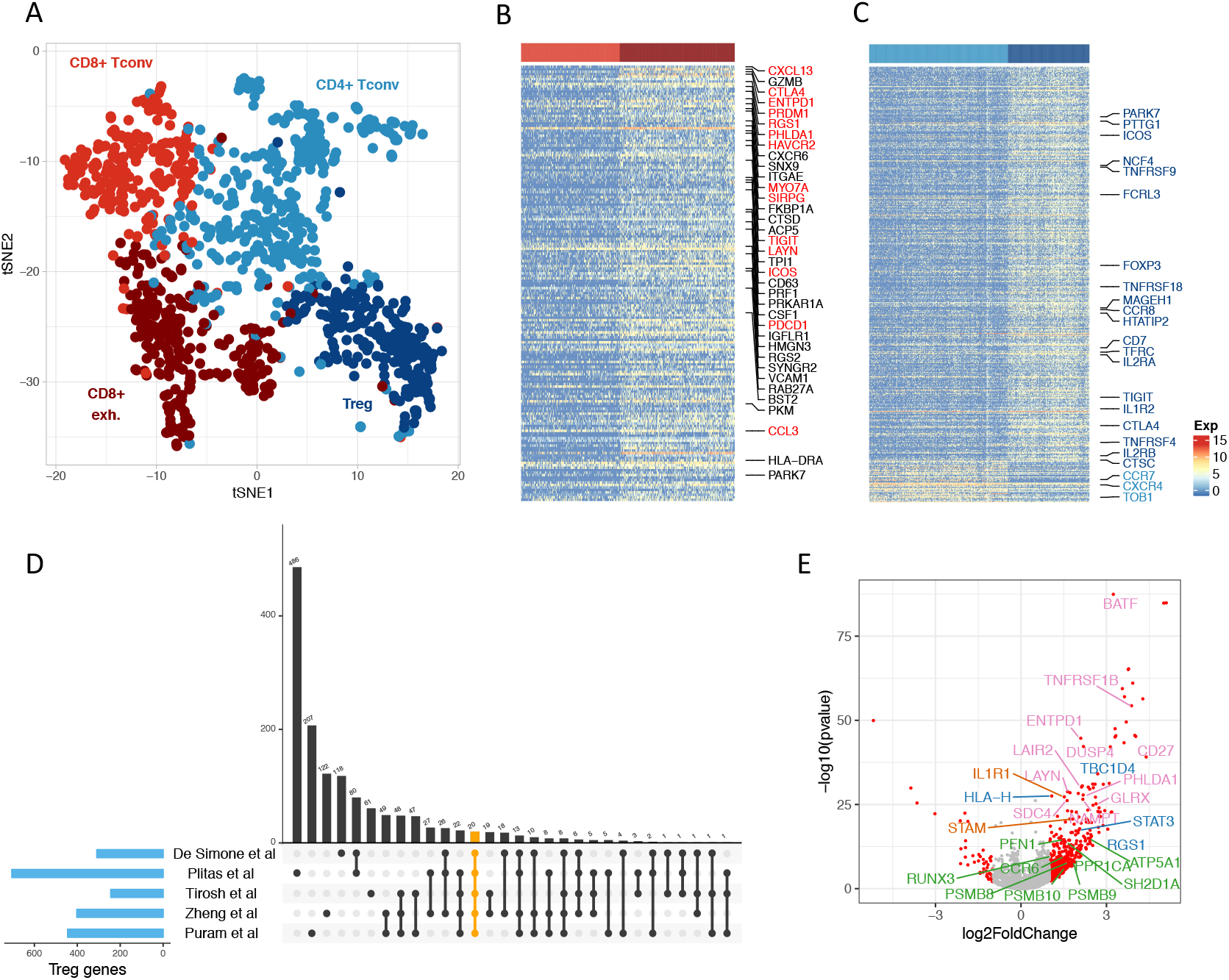
Deconvolution of T cell subtypes. (A) 2D t-sne projection of T cells. T cell subtypes identified by clustering analysis are annotated and marked by color codes. (B) Heatmap of genes significantly expressed in exhausted CD8^+^ T cells comparing to conventional CD8^+^ T cells (adjusted p-value < 0.05, log2fold-change > 1). Genes also reported by a previous study are labeled on left, of which the known exhaustion markers are labeled in red text. Cell types are indicated by the colored bar at top. (C) Heatmap of genes differentially expressed in Tregs comparing with conventional CD4^+^ T cells (adjusted p-value < 0.05, |log2fold-change| > 1). Selective Treg genes are labeled in dark blue and known markers for conventional CD4^+^ T cells are labeled in light blue. (D) Intersections between Treg genes identified in (C) using Puram et al. HNSCC data with those from previous studies. Numbers of genes in each intersection are shown as the bars at the top panel. Studies involved in each intersection are linked together by lines at the bottom panel. The 20 genes shared by all five studies are highlighted in orange and labeled in (C). (E) Volcano plot of genes differentially expressed in Tregs vs. conventional CD4^+^ T cells. Unique genes found by this study are labeled in green. Those identified once (blue), twice (red), and three times (pink) previously are also labeled.

### Evaluation of prediction performance of reference GEPs

For each cell type identified from previous steps, we established cell-type-specific reference GEP matrix by the mean expression values of selected genes. We use C1 to denote the curated gene list from previous literatures which are used in ssGSEA (Table S2), C2 to denote marker genes selected from the DE analysis described above (Table S4), T1 to denote the marker genes selected from DE analyses for separating T cell subtypes (Table S3), and M1 to denote marker genes selected from DE analyses for separating tumor and non-tumor cells. In our analysis, we constructed reference GEP matrices by taking the mean from the following ensemble gene lists: (1) LM22, (2) C1, (3) C2, (4) LM22+C1, (5) LM22+C1+T1, (6) LM22+C1+T1+M1, and (7) LM22+C1+C2+T1+M1. As presented in Figure S5, we evaluated the prediction performance of CIBERSORT using these GEPs in terms of correlation between predicted abundance and the true abundance in the simulated bulk tumor (through pooling all cells in one patient, see Methods). We observed that all of these reference GEPs achieved promising prediction accuracies (r>0.9). This result indicates that existing marker genes provides saturated signatures if forming GEPs on right cell groups. Therefore, we will focus on the evaluation of the LM22+C1 gene panel because of it has a moderate number of genes and all genes included are well studied. All reference GEPs matrices used in this study are provided in Supplementary Table S5.

Scatterplots in Figure 3A demonstrate strong correlations between true cell proportions and predicted cell proportions based on GEP curated form LM22+C1 scRNA-seq data, where each point represents a simulated bulk sample. Figure 3B further compares the cell abundance estimation accuracy (correlation) for the reference GEP included in CIBERSORT and the reference GEP trained based on the LM22+C1 scRNA-seq panel. Our method shows better prediction performance in all case for cell types that CIBERSORT can provide estimation, especially in estimating CD8 T cells. We further gauged the estimated cell proportion from CIBERSROT by taking into account the fact that the original GEP only include reference for immune cells. Such adjustment was made by assuming that tumor cell (purity) and stromal cell proportion were known so that a relative abundance on each remaining cell types can be calculated. Even with this unrealistic scenario, the prediction performance based on the adjusted proportion was still inferior to the scRNA-seq trained GEP in all cases. But we did observe that CIBERSORT estimation on macrophages and dendritic cells was greatly improved with this adjustment (Figure S6). To test the robustness of the GEP panel to the cell components, we re-run the devolution analysis on all simulated samples using the leave-one-out GEP, i.e. each time we remove one cell-type-specific vector from the GEP matrix. As shown in Figure S7, the high prediction accuracy was maintained in most scenarios, and only the estimations for fibroblasts and malignant cells were detectably impacted by the leave-one-out GEP.

**Figure 3.**
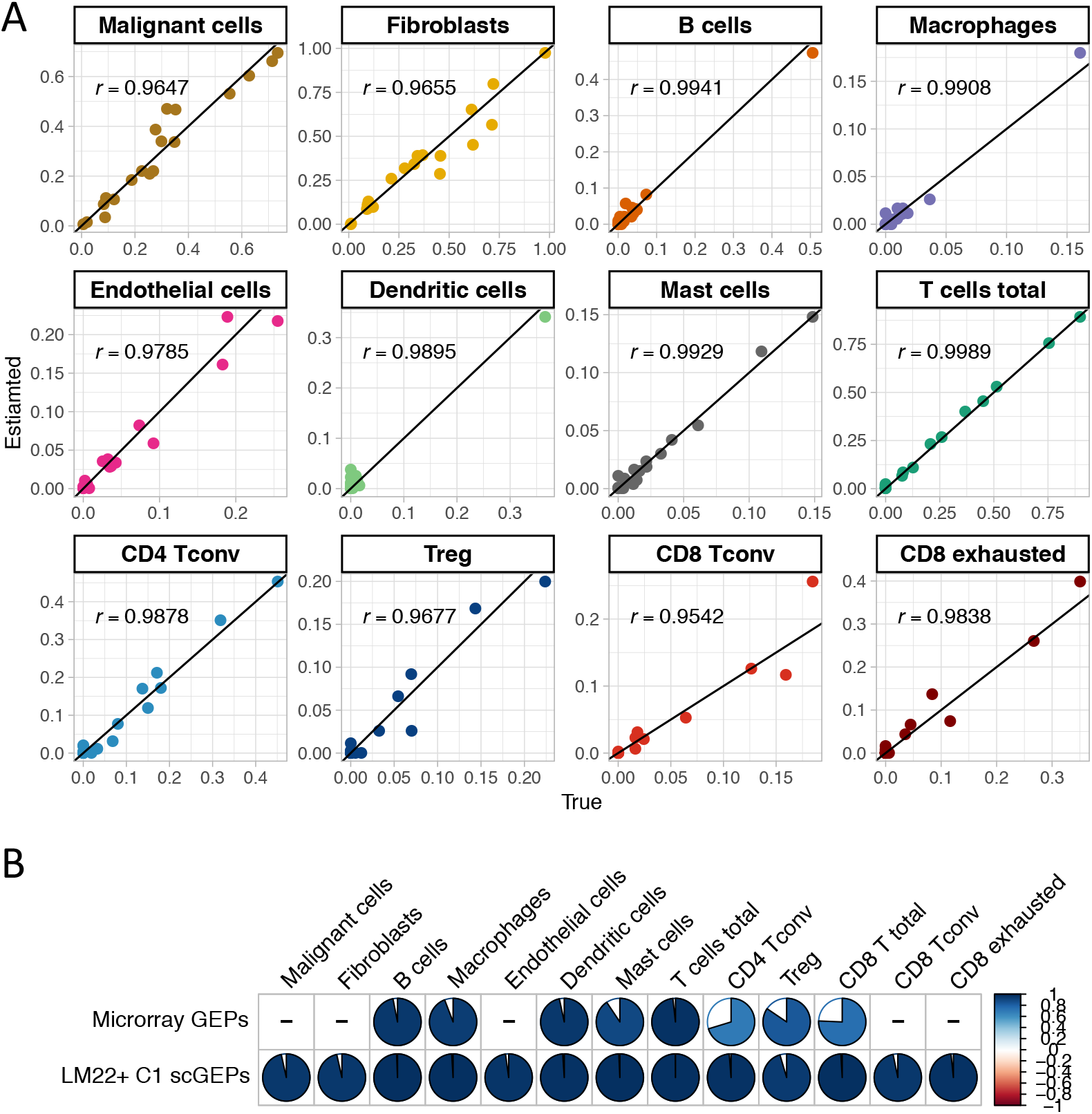
Estimation accuracy of cellular compositions using LM22^+^C1 scGEPs. (A) Scatter plots of the estimated and true cell proportions for the 20 simulated bulk tumor samples. Each dot represents one sample and r denotes the Pearson’s correlation coefficient. (B) Comparing the estimation accuracy between LM22^+^C1 scGEPs and CIBERSORT microarray GEPs. The pie charts show the Pearson’s correlation coefficient of true proportion and proportion estimated using CIBERSORT microarray GEPs (top), and LM22^+^C1 scGEPs (bottom). The missing cell types in CIBERSORT microarray GEPs are denoted by dash. T cell composition was calculated taking sum of the four T cell subtypes.

Although C2 and T1 gene sets (determined based on DE tests) did not provide additional information as a gene panel in constructing GEP, they provide a new alternative cell-type-specific biomarker for future studies. As shown in violin plots (Figure S14), these markers are exclusively over-expressed in cell types that they are representing, indicating their validity as independent surrogate biomarkers. A total of 182 genes were found overlapping between groups C2+T1 and LM22+C1. Expressions of these genes for each single cell were plotted in Figure S15, demonstrating their ability as biomarker panel alone to separate major cell types but not T cell subtypes.

Finally, as a supportive validation, we tested the proposed scGEP on TCGA HNSCC tumor samples and compared with results generated from similar methods developed for bulk tumor deconvolution. Figure S8 compared the tumor purity estimates with three other methods: ABSOLUTE [21], ESTIMATE [22] and CPE [23]. These methods are based on WES, RNAseq and a consensus score based on all molecular data. Our method showed the best correlation with the estimation from ESTIMATE in terms of purity estimation. Further, we compared the Immune and Stromal score predicted by ESTIMATE with the absolute proportion estimates from the scGEP-based method. As shown in Figure S9, the analysis showed a good agreement between two methods. We also compared the estimated total immune cell proportions and total T cell proportions between HPV positive and HPV negative cancer patients. As expected, tumors from HPV positive patients showed higher infiltration of immune cells and T cells (Figure S10). Abundance of tumor infiltrating CD8 and total immune cells were also found associated with survival outcomes in TCGA HNSCC patients (Figure S11).

## Discussion

scRNA-seq provides high resolution data to study cell heterogeneity, and provides new chance to understand the dynamic ecosystem comprising tumor cells, fibroblasts, and immune cells. Nevertheless, gene expression data from bulk tumors is indispensable and still dominates the clinical and translational settings. In this study we developed a pipeline to construct the reference gene expression profile matrix based on scRNA-seq data (scGEP), and assessed its performance in estimating cancer and immune cell compositions from bulk tumor gene expression data. By combining gene expression profiles of major cancer and immune cell types in HNSCC established from a high-quality single cell data, our approach overcomes a key shortcoming of most existing studies that relied on limited source of FACS-purified cell populations for the reference signature gene matrix. As noted in previous studies, PBMC-based GEP is also insufficient to provide accurate estimate on bulk tumor samples. The scGEP matrix derived from our analysis provides a new resource for future endeavors in analyzing expression data in head and neck cancers. The estimation on tumor purity will be greatly improved with the tailored reference signature for HNSCC malignant cells. Importantly, more accurate estimation on cancer cells partly contributes to better estimation on the relative abundance of immune cells. We validated results by using in silico pooled bulk tumor samples, and also showed that single-cell-derived signatures provides the ability to separate T cell subtypes. The finer and more accurate tumor immune profiling of HNSCC samples will help reveal more prognostic biomarkers with implications for immunotherapy. Furthermore, because immune cell share very similar expression profiles across cancer types, in theory the reference matrix can be broadly employed to other solid tumors, but it will only provide relative abundance for immune cell types. With the increased availability of single-cell data in cancers such as melanoma and lung cancers, an ideal scGEP matrix should be generated based on the same tumor type using the proposed pipeline.

The key step in constructing scGEP matrix involves accurately identifying cells of the same types or subtypes from heterogeneous populations, which is the in-silico equivalent of isolating cells using physical sorting methods. Compared to traditional sorting methods such as FACS, in-silco methods are less time consuming, less laborious, and more cost effective. Cell type determination at cellular level have benefited greatly from specialized clustering methods developed for scRNA-seq [8, 14, 24–26]. While there are more advanced approaches including deep learning [27, 28] have been proposed in recent years, fully automated decomposition of cell types is still a challenging problem. Part of the difficulty arises from the fact that each tumor includes a large variety of malignant and nonmalignant cells at different stages. The cellular mixing component and proportions even with the same section of a tumor can be very different if sampled under different time or conditions, e.g., before or after treatment. In addition, due to the limitations of the scRNA-seq technology itself, single cell gene expression data are often very noisy. And hence cells of the same type can end up in different clusters, and cells of different types can be in the same cluster due to unknown technology batch effects. Therefore, it is important to carefully curate and select high-quality cell clusters before calculating cell-type-specific reference matrix. In this study, we adopted an adaptive divide-and-conquer scheme to identify all major cell types in HNSCC tumor tissues, starting from the easiest split of cancer vs. non-cancer cells to the most challenging T cell subtype separation. In every step of the process, cell types are inferred based on both the results from the unsupervised clustering analysis and the expression status of existing marker genes. Any prior knowledge about the cellular component of the studied tumor type also helps in assigning a cell cluster to a cell types. In later stages of the analysis where cell subtypes are getting harder to distinguish, multiple settings or even multiple methods of clustering analysis need to tired. This process cannot be automated due to the need to visually inspect the clustering results in each step, but can achieve the best possible results for cell mixture deconvolution. The adaptive method was also based on a key assumption and it was further demonstrated in our clustering analysis: despite the significant heterogeneity of malignant cells across tumors, cells from the same immune cell types can be clustered together due to their relatively similar gene expression profiles.

It is important to highlight main advantage of using the data from Puram et al. as training dataset: it by far contains the largest collection of single cells from solid tumors (in terms of patient number and cell number) from a single study. The above-described property allows us to calculate the composite reference GEP not only from pooled cells from different tumor samples, but also from different scRNA-seq experiments. A caveat is that data pooled across studies involves more complicated batch effects and it is by now generally accepted that correcting for the batch effect in RNAseq data across experiments is technically challenging. It is interesting to note, though, that some recently proposed ideas for batch effect correction with scRNA-seq data are based on consensus clustering, which leverages the same philosophy mentioned above by projecting more homogenous immune cells into the same cluster. As pointed out in the original analysis, some apparent batch effect observed may be linked to the enzyme used for reverse transcription in the scRNA experiments. We further investigated the factor of enzyme usage in the adaptive clustering scheme and found that it explains well the sub-clusters observed in Fibroblast cell populations (Figure S12), but had limited impact on other cell types (Figure S13).

One notable observation in our simulation studies is that the GEP calculated based on existing marker gene panel (LM22+C1) can provide as accurate a predictive capability as the genes selected only from the differential expression of single cell populations, although they are overlapped in many genes. We conclude that the prediction performance of GEP is more sensitive to the cell populations purified for a particular cell type than the marker gene panel. Nevertheless, the newly discovered genes from the scRNA-seq data and their underlying pathway warrant further validations as potential biomarkers, especially those genes that are differentially expressed between T subtypes.

Although we have only tested support vector regression method for cell mixture estimation, the HNSCC single cell sequencing data curated from this study provides a useful source for the assessment of accuracy of newly developed deconvolution methods. For example, the core SVR algorithm implemented in CIBESRORT only uses a single kernel under the fixed default parameter setting. The prediction performance might be improved through searching for an optimal kernel or using the state-of-the-art multiple kernel learning technologies [24]. Currently there is a lack of suitable benchmark dataset that allows a fair and systematic evaluation of methods for estimating cell mixtures in solid tumors. Weak correlations were often found between molecular-data-based estimations and pathology based methods such as IHC and H&E images [29]. This is partially due to the fact that each of these assays was carried out using input materials from different parts of a tumor. Because all cell proportions are known, the in silico pooled bulk tumor data from individual cells provides a more accurate reference at almost zero cost. Plus the composite cells from a single tumor could better mimic the real case scenario than creating bulk expression dataset through conducting RNA-seq on randomly mixed cells. For head and neck cancer per se, the scRNA-seq data from Puram study provide an ideal source for both training and validation purposes because the studied tumors have (1) uniformly varied tumor purity, and (2) it provides reference for subpopulations such as exhaustive CD8^+^ T cell that were not present in previous scRNA-seq experiments on melanoma and lung cancers. A limitation of the in silico method is that the cell size factor has not been taken into account. As cell types of different size have different amount of RNA yield, it is of interest for future research to be able to adjust for the cell size factor so that the estimated relative abundance will be closer to absolute cell proportions.

The key idea proposed in this work is most similar to a previous study conducted by Schelker et al. [7] which focused on scRNA-seq data from melanoma and PBMC. The two main differences between the two works are (1) the melanoma data used by Schelker et al. only provided sufficient information for distinguish nine major cell types and three T cell subtypes, whereas the HNSCC data we studied was able to further separate exhaustive CD8 T cells and provide corresponding reference GEP; (2) In our method, we used both marker gene information and a global ssGSEA scores to determine cell types from adaptive clustering analysis. We believe that more studies along this line will be conducted to generate more accurate cancer-type-specific and T-cell-subtype-specific reference GEP. Finally, we believe that apart from looking for reference profiles based on gene expression, the same approach can be extended in future search to identify reference DNA methylation profiles (DMP). DMP will be a promising new resource for tumor composition deconvolution because Alternations at DNA methylation level are deemed to be more stable than the gene expression level. But the single-cell DNA methylation analysis, such as bisulfite sequencing, is still in an experimental phase.

## Conclusions

## List of abbreviations

scRNA-seq: single-cell RNA sequencing
TILS: tumor-infiltrating lymphocytes
IHC: immunohistochemical
t-SNE: t-Distributed Stochastic Neighbor Embedding
HNSCC: head and neck squamous cell carcinoma
GEP: gene expression profiles
scGEPs: single-cell gene expression profiles
TME: tumor microenvironment
CD4^+^ Tconv: conventional CD4^+^ T cells
CD8^+^ Tconv: conventional CD8^+^ T cells
CD8^+^ exhausted: exhausted CD8^+^ T cells
TPM: transcripts per million
ssGSEA: the single-sample Gene Set Enrichment Analysis
v-SVR: v-support vector regression

## Declarations

### Ethics approval and consent to participate

Not applicable.

### Consent for publication

Not applicable.

### Availability of data and materials

The HNSCC scRNA_seq data used in this study was retrieved from Puram et al [1].

### Competing interests

The authors declare that they have no competing interests.

### Funding

This work was supported in part by Institutional Research Grant number 14-189-19 from the American Cancer Society, and a Department Pilot Project Award from Moffitt Cancer Center. The funders had no role in study design, data collection and analysis, decision to publish, or preparation of the manuscript.

### Authors’ contributions

All authors read and approved the final manuscript. XY, YAC, JRC, CHC and XW conceived the study, XY, YAC and XW designed the algorithm. XY and XW performed the analyses, interpreted the results and wrote the manuscript.

## Acknowledgments

The authors would like to thank Colleagues at Department of Biostatistics and Bioinformatics at Moffitt Cancer Center for providing feedback.

## References

1. Puram SV, Tirosh I, Parikh AS, Patel AP, Yizhak K, Gillespie S, Rodman C, Luo CL, Mroz EA, Emerick KS et al: Single-Cell Transcriptomic Analysis of Primary and Metastatic Tumor Ecosystems in Head and Neck Cancer. Cell 2017, 171(7): 1611–1624.e1624.

2. Tanay A, Regev A: Scaling single-cell genomics from phenomenology to mechanism. Nature 2017, 541(7637):331–338.

3. Tirosh I, Izar B, Prakadan SM, Wadsworth MH, 2nd, Treacy D, Trombetta JJ, Rotem A, Rodman C, Lian C, Murphy G et al: Dissecting the multicellular ecosystem of metastatic melanoma by single-cell RNA-seq. Science (New York, NY) 2016, 352(6282): 189–196.

4. Wagner GP, Kin K, Lynch VJ: Measurement of mRNA abundance using RNA-seq data: RPKM measure is inconsistent among samples. Theory in Biosciences 2012, 131(4):281–285.

5. Barbie DA, Tamayo P, Boehm JS, Kim SY, Moody SE, Dunn IF, Schinzel AC, Sandy P, Meylan E, Scholl C et al: Systematic RNA interference reveals that oncogenic KRAS-driven cancers require TBK1. Nature 2009, 462(7269):108–112.

6. Şenbabaoğlu Y, Gejman RS, Winer AG, Liu M, Van Allen EM, de Velasco G, Miao D, Ostrovnaya I, Drill E, Luna A et al: Tumor immune microenvironment characterization in clear cell renal cell carcinoma identifies prognostic and immunotherapeutically relevant messenger RNA signatures. Genome biology 2016, 17(1):231–231.

7. Schelker M, Feau S, Du J, Ranu N, Klipp E, MacBeath G, Schoeberl B, Raue A: Estimation of immune cell content in tumour tissue using single-cell RNA-seq data. Nature communications 2017, 8(1):2032–2032.

8. Kiselev VY, Kirschner K, Schaub MT, Andrews T, Yiu A, Chandra T, Natarajan KN, Reik W, Barahona M, Green AR et al: SC3: consensus clustering of single-cell RNA-seq data. Nature methods 2017, 14(5):483–486.

9. Ritchie ME, Phipson B, Wu D, Hu Y, Law CW, Shi W, Smyth GK: limma powers differential expression analyses for RNA-sequencing and microarray studies. Nucleic acids research 2015, 43(7):e47–e47.

10. Schölkopf B, Smola AJ, Williamson RC, Bartlett PL: New Support Vector Algorithms. Neural Comput 2000, 12(5):1207–1245.

11. Newman AM, Liu CL, Green MR, Gentles AJ, Feng W, Xu Y, Hoang CD, Diehn M, Alizadeh AA: Robust enumeration of cell subsets from tissue expression profiles. Nature methods 2015, 12(5):453–457.

12. Mandal R, Şenbabaoğlu Y, Desrichard A, Havel JJ, Dalin MG, Riaz N, Lee K-W, Ganly I, Hakimi AA, Chan TA et al: The head and neck cancer immune landscape and its immunotherapeutic implications. JCI insight 2016, 1(17):e89829–e89829.

13. Zheng C, Zheng L, Yoo J-K, Guo H, Zhang Y, Guo X, Kang B, Hu R, Huang JY, Zhang Q et al: Landscape of Infiltrating T Cells in Liver Cancer Revealed by Single-Cell Sequencing. Cell 2017, 169(7):1342–1356.e1316.

14. Duò A, Robinson MD, Soneson C: A systematic performance evaluation of clustering methods for single-cell RNA-seq data. F1000Research 2018, 7:1141–1141.

15. Plitas G, Konopacki C, Wu K, Bos PD, Morrow M, Putintseva EV, Chudakov DM, Rudensky AY: Regulatory T Cells Exhibit Distinct Features in Human Breast Cancer. Immunity 2016, 45(5):1122–1134.

16. De Simone M, Arrigoni A, Rossetti G, Gruarin P, Ranzani V, Politano C, Bonnal RJP, Provasi E, Sarnicola ML, Panzeri I et al: Transcriptional Landscape of Human Tissue Lymphocytes Unveils Uniqueness of Tumor-Infiltrating T Regulatory Cells. Immunity 2016, 45(5):1135–1147.

17. Sugai M, Aoki K, Osato M, Nambu Y, Ito K, Taketo MM, Shimizu A: Runx3 Is Required for Full Activation of Regulatory T Cells To Prevent Colitis-Associated Tumor Formation. The Journal of Immunology 2011, 186(11):6515–6520.

18. Yamazaki T, Yang XO, Chung Y, Fukunaga A, Nurieva R, Pappu B, Martin-Orozco N, Kang HS, Ma L, Panopoulos AD et al: CCR6 regulates the migration of inflammatory and regulatory T cells. Journal of immunology (Baltimore, Md: 1950) 2008, 181(12):8391–8401.

19. Ruszkowski J, Lisowska KA, Pindel M, Heleniak Z, Dębska-Ślizień A, Witkowski JM: T cells in IgA nephropathy: role in pathogenesis, clinical significance and potential therapeutic target. Clinical and experimental nephrology 2019, 23(3):291–303.

20. Li Z, Li D, Tsun A, Li B: FOXP3+ regulatory T cells and their functional regulation. Cellular & molecular immunology 2015, 12(5):558–565.

21. Carter SL, Cibulskis K, Helman E, McKenna A, Shen H, Zack T, Laird PW, Onofrio RC, Winckler W, Weir BA et al: Absolute quantification of somatic DNA alterations in human cancer. Nature Biotechnology 2012, 30:413.

22. Yoshihara K, Shahmoradgoli M, Martínez E, Vegesna R, Kim H, Torres-Garcia W, Treviño V, Shen H, Laird PW, Levine DA et al: Inferring tumour purity and stromal and immune cell admixture from expression data. Nature Communications 2013, 4:2612.

23. Aran D, Sirota M, Butte AJ: Systematic pan-cancer analysis of tumour purity. Nature communications 2015, 6:8971–8971.

24. Wang B, Ramazzotti D, De Sano L, Zhu J, Pierson E, Batzoglou S: SIMLR: A Tool for Large-Scale Genomic Analyses by Multi-Kernel Learning. PROTEOMICS 2018, 18(2):1700232.

25. Butler A, Hoffman P, Smibert P, Papalexi E, Satija R: Integrating single-cell transcriptomic data across different conditions, technologies, and species. Nature Biotechnology 2018, 36:411.

26. Langfelder P, Horvath S: WGCNA: an R package for weighted correlation network analysis. BMC bioinformatics 2008, 9:559–559.

27. Lopez R, Regier J, Cole MB, Jordan MI, Yosef N: Deep generative modeling for single-cell transcriptomics. Nature Methods 2018, 15(12):1053–1058.

28. Cho H, Berger B, Peng J: Generalizable and Scalable Visualization of Single-Cell Data Using Neural Networks. Cell Systems 2018, 7(2):185–191.e184.

29. Saltz J, Gupta R, Hou L, Kurc T, Singh P, Nguyen V, Samaras D, Shroyer KR, Zhao T, Batiste R et al: Spatial Organization and Molecular Correlation of Tumor-Infiltrating Lymphocytes Using Deep Learning on Pathology Images. Cell reports 2018, 23(1):181–193.e187.

