## Supplemental Figures for "Estimation of immune cell content in tumor using single-cell RNA-seq reference data"

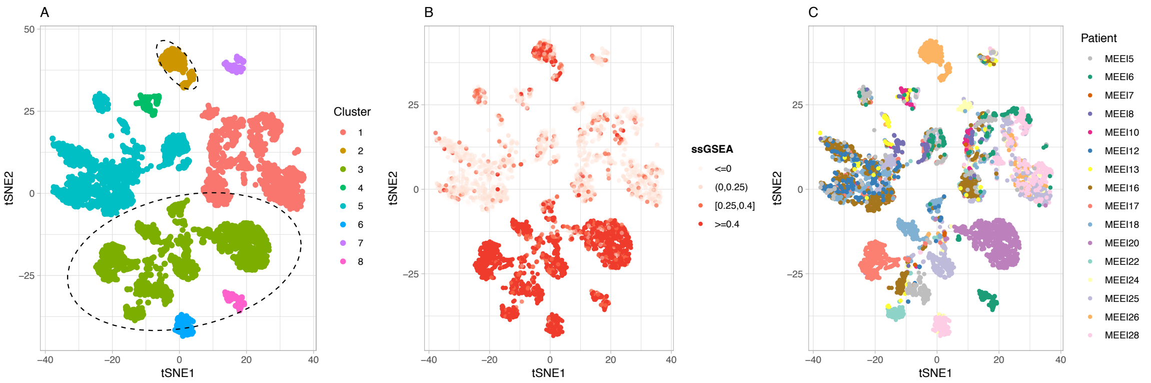


**Figure S1. Separating malignant cells from immune and stromal cells.**
(A) The same 2D t-sne projection of all 5,712 single cells as in Figure 1A. Clusters identified by t-SNE and DBscan clustering (with parameters eps=5 and minPts =5) are shown in different colors. Clusters considered as malignant cells are circled.
(B) The same 2D t-sne projection as in (A) with cells colored by ssGSEA score calculated using HNSCC malignant cell markers in Supplementary Table S2.
(C) The same 2D t-sne projection as in (A) and (B) with cells colored by patient origins.

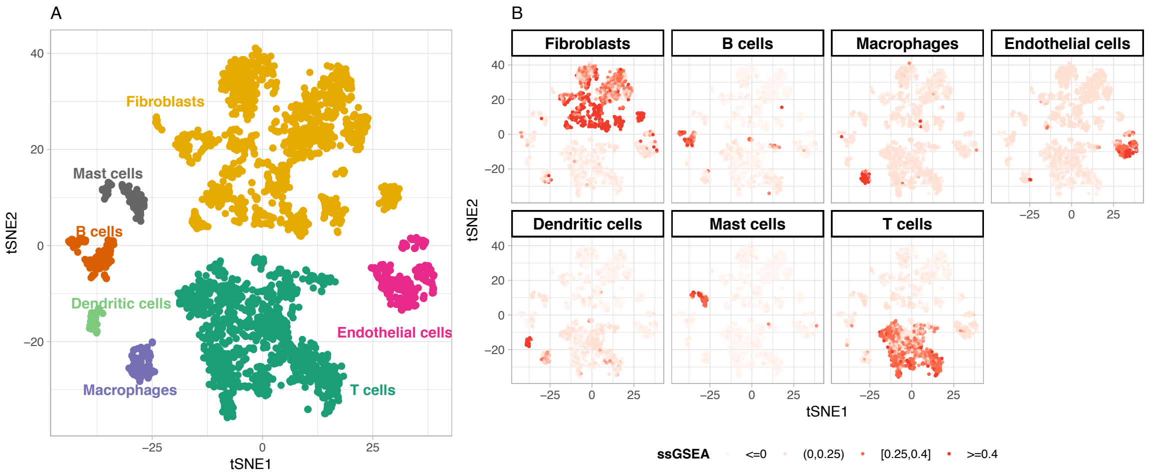


**Figure S2. Identifying major immune and stromal cell-type clusters.**
(A) 2D t-sne projection of the 3,259 non-malignant single cells. Clusters identified by t-SNE and DBscan clustering (with parameters eps=6 and minPts =15) are associated with cell types (colors).
(B) The same 2D t-sne projection as in (A) with cells colored by ssGSEA scores calculated using signature genes for fibroblasts, B cells, macropahges, endothelia cells, dendritic cells, mast cells, and T cells (Supplementary Table S2), respectively.

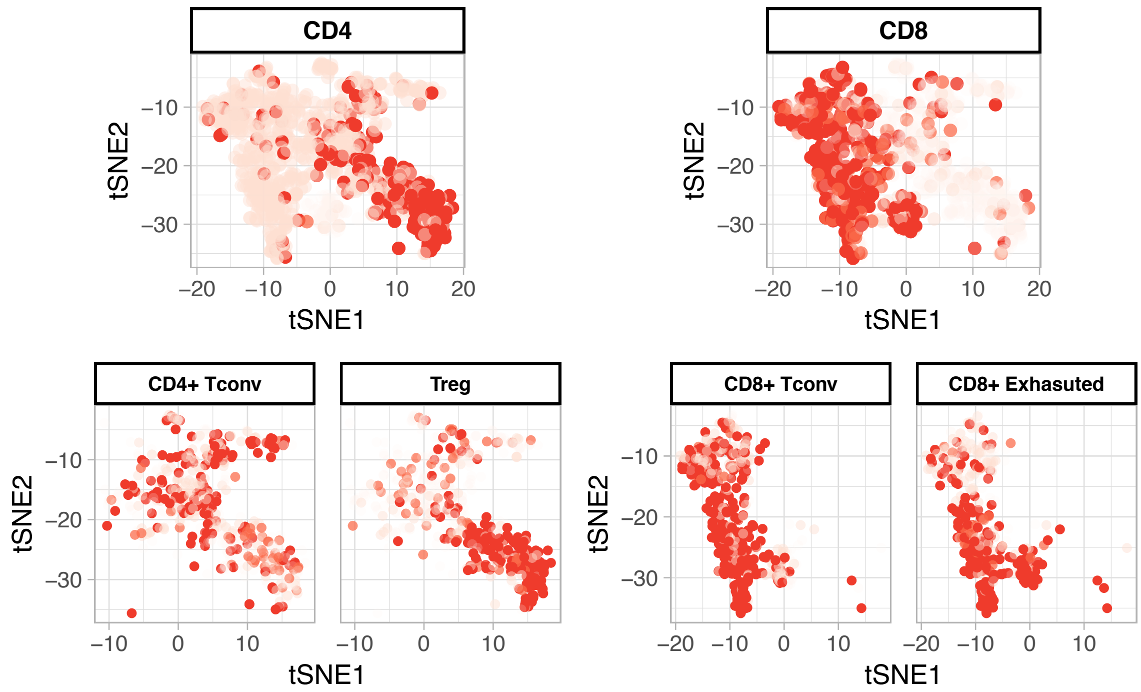


**Figure S3. ssGSEA scores for CD4$^+$ and CD8$^+$ T cell subtypes.**
Showing is the same 2D t-sne projection of T cells as in Figure 2A. The cells are colored by ssGSEA scores calculated using signature genes for CD4$^+$, CD8$^+$, conventional CD4$^+$, Treg, conventional CD8$^+$, and exhausted CD8$^+$ T cells (Supplementary Table S2).

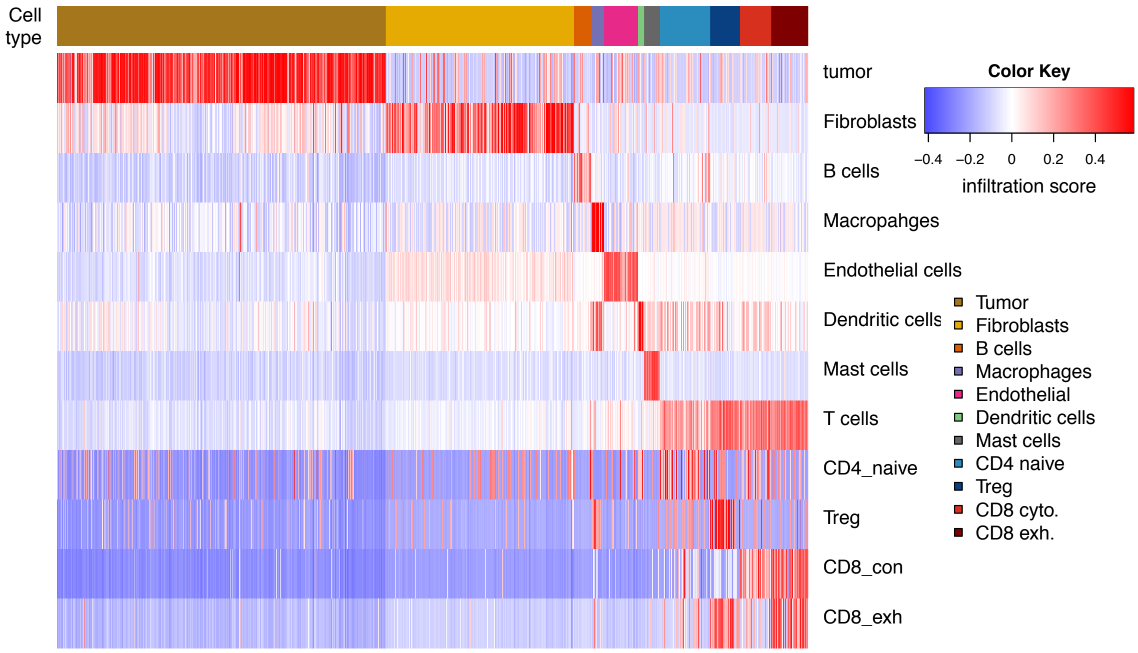


**Figure S4. Heatmap of the ssGSEA scores of 12 cell types across all single cells.** Columns are single cells and the top bar indicates the final assignment of cell types. Rows are the ssGESA score calculated for single cells using signature genes of a specific cell type.

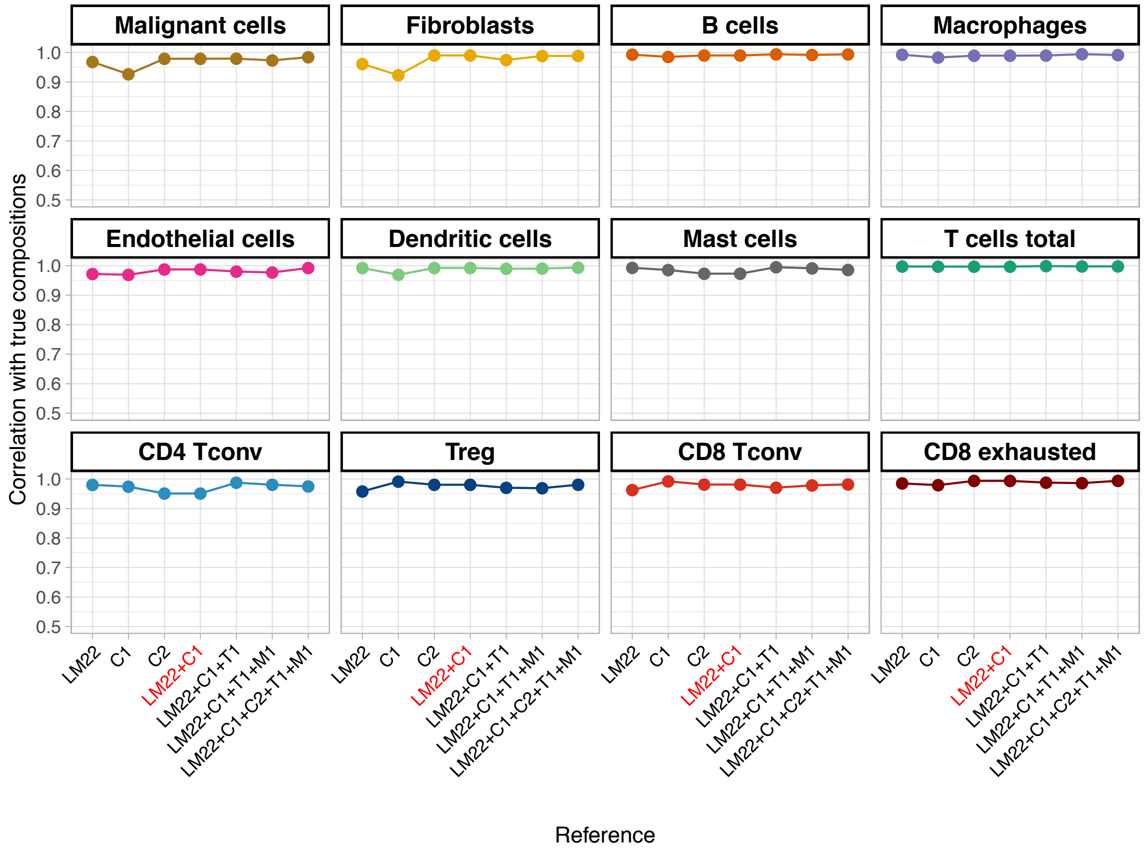


**Figure S5. Estimation accuracy of cellular compositions using different scGEPs.** Estimation accuracy is measured as the Pearson's correlation coefficient of true cell proportion vs. cell proportion estimated using scGEPs. For each cell type, the accuracy is calculated for 7 sets of scGEPs: (1) LM22, (2) C1, (3) C2, (4) LM22+C1, (5) LM22+C1+T1, (6) LM22+C1+T1+M1, and (7) LM22+C1+C2+T1+M1.

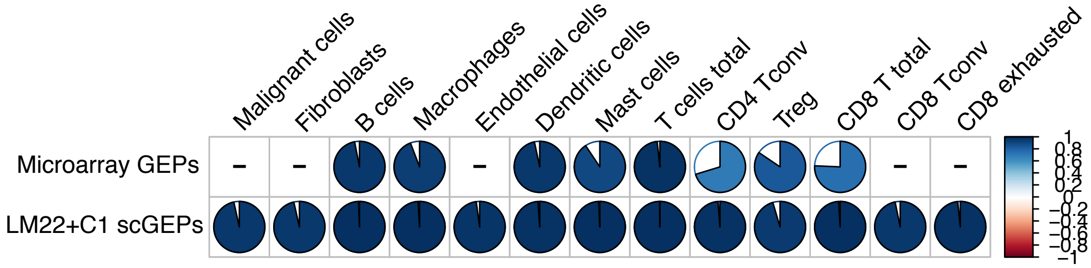


**Figure S6. Comparing the estimation accuracy between LM22+C1 scGEPs and CIBERSORT microarray GEPs, based on adjusted proportion. Related to Figure 3B**. Bottom row is the same as Figure 3B, which shows the Pearson's correlation coefficient of true proportion vs. proportion estimated by LM22+C1 scGEPs. Top row shows the Pearson's correlation coefficient of true proportion vs. adjusted proportion estimated by CIBERSORT microarray GEPs. For a specific cell type, the adjusted proportion = estimated proportion / (1- true malignant cell proportion – true stromal cell proportion).

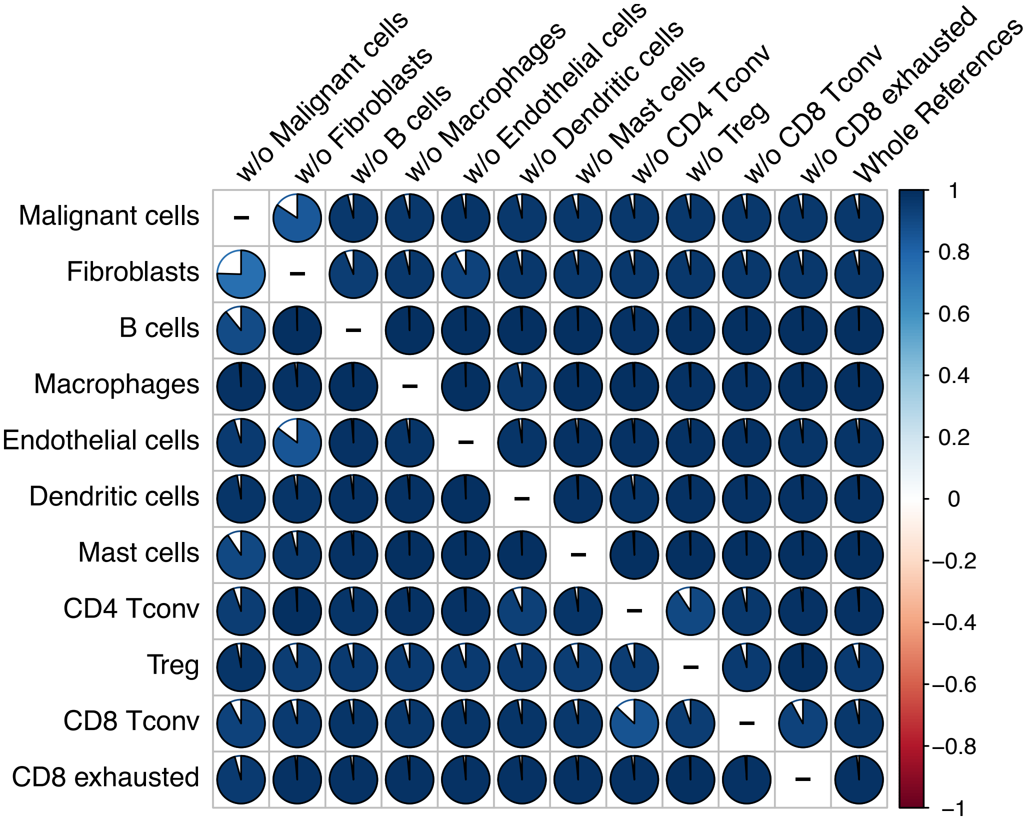


**Figure S7. The impact of missing cell types in scGEPs.**
The pie charts show the Pearson's correlation coefficient of true proportion vs. proportion estimated using scGEPs matrix with one cell type removed. Rows are cell type, and columns are leave-one-out scGEPs.

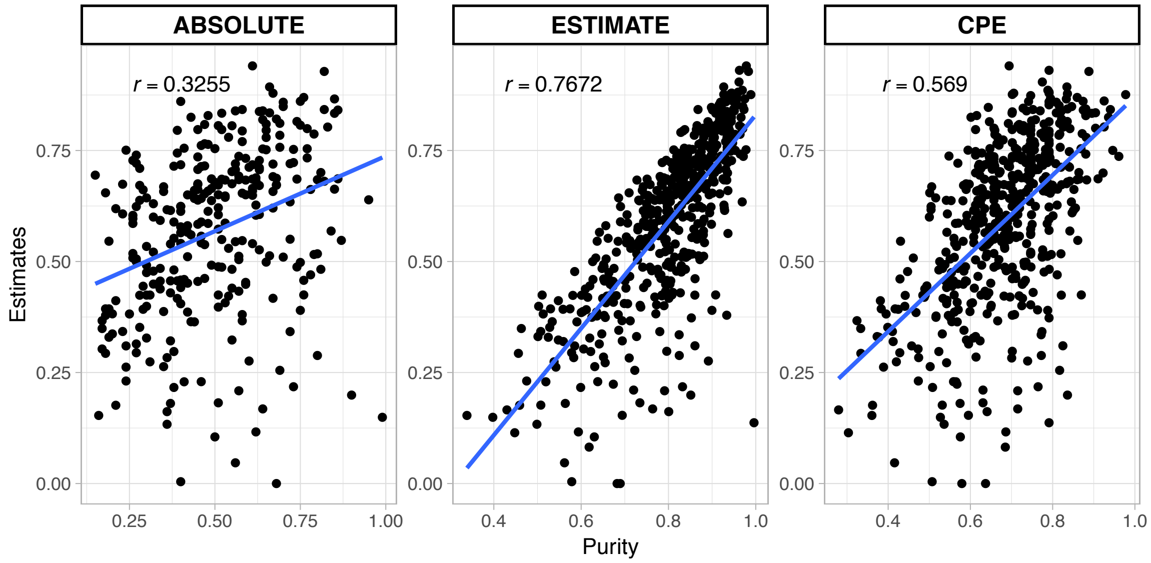


**Figure S8. Malignant cell proportion estimated by scGEPs vs. tumor purity in TCGA HNSCC bulk tumor RNA-seq data**

Scatter plots of malignant cell proportion estimated for TCGA HNSCC tumor RNA-seq data using scGEPs vs. tumor purity predicted by three methods: ABSOLUTE, ESTIMATE, and CPE. Each dot represents a sample and r denotes the Pearson's correlation coefficient.

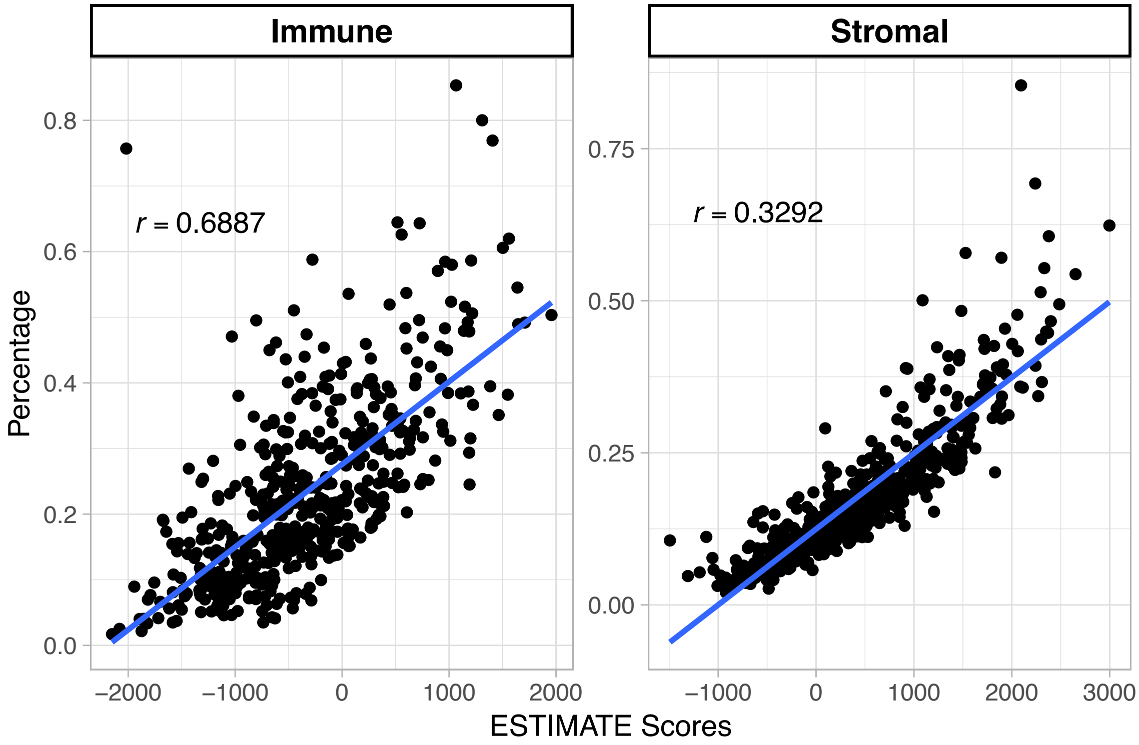


**Figure S9. Comparing the estimated immune and stromal proportions with ESTIMATE scores in TCGA HNSCC bulk tumor RNA-seq data.**
Scatter plots of immune and stromal cell proportions estimated using scGEPs vs. Immune and Stromal score predicted by ESTIMATE. Each dot represents a sample and r denotes the Pearson's correlation coefficient.

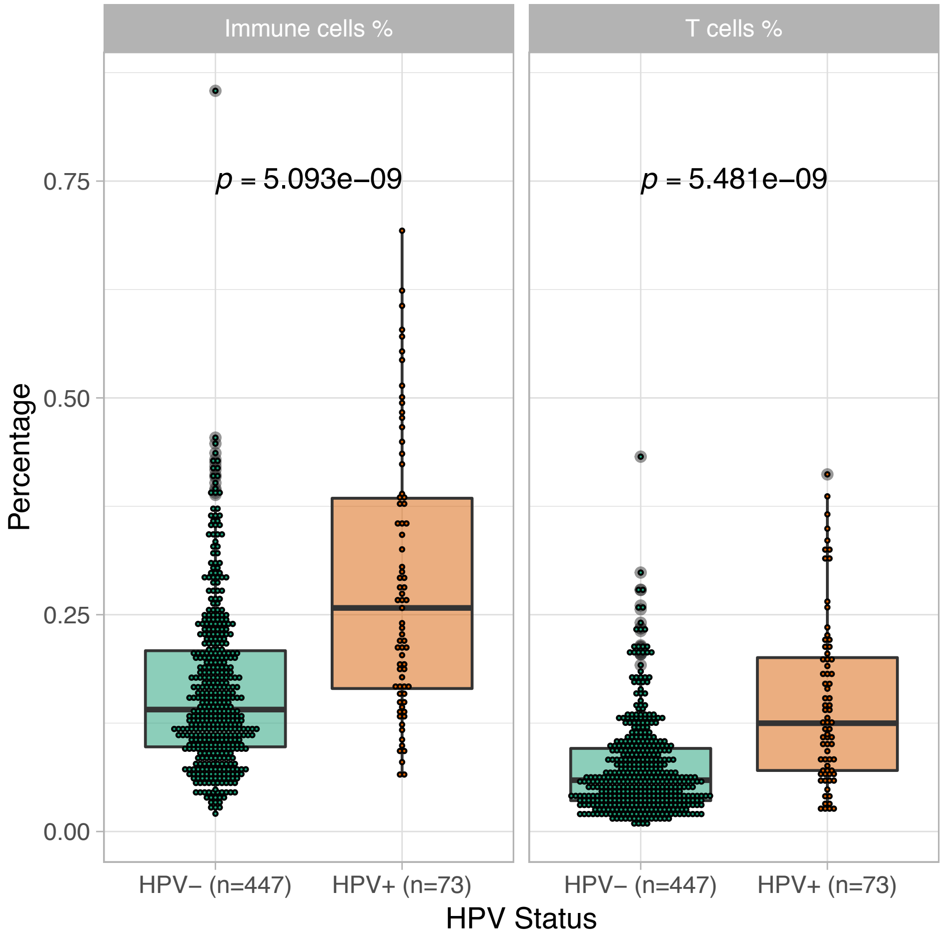


**Figure S10. Association between immune/T cell proportion and HPV status in TCGA HNSCC**

Boxplots shows the immune cell and T cell proportion estimated by scGEPs in $HPV^+$ and $HPV^-$ TCGA HNSCC patients respectively. P-value of Mann-Whitney-Wilcoxon Test comparing proportion $HPV^+$ vs. $HPV^-$ patients is also shown.

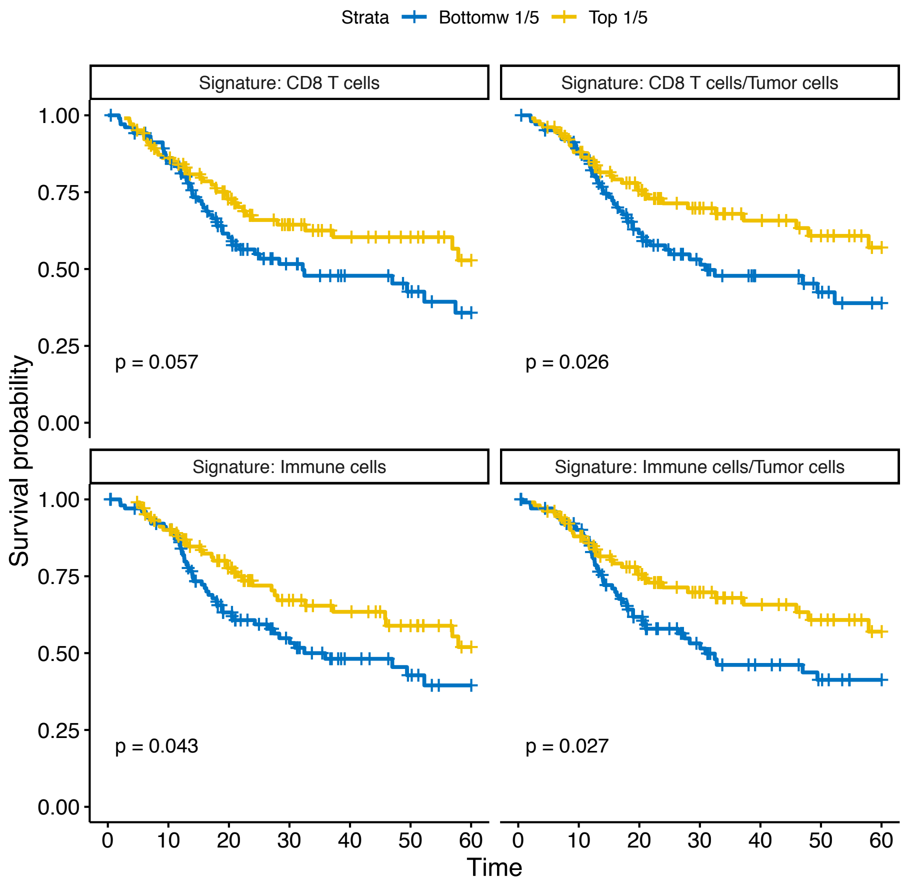


**Figure S11. Association of estimated cellular compositions with overall survival in TCGA HNSCC patients.**

TCGA HNSCC patients are categorized into top 20% and bottom 20% based on 4 signatures calculated from estimated cellular composition: (1) CD8 T cell proportion, (2) the ratio of CD8 T cell proportion / tumor cell proportion, (3) immune cell proportion, or (4) the ratio of immune cell proportion / tumor cell proportion. Sixth overall month survival curves of these two groups of patients are compared using log-rank test and the p-values are shown in plot.

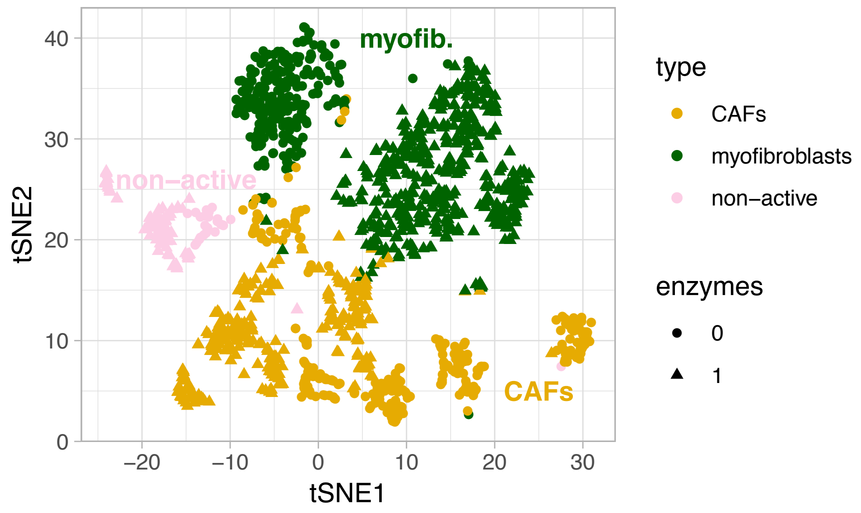


**Figure S12. Identification of fibroblast cell subtypes.**
2D t-sne projection of the 1,451 fibroblasts single cells with cells. Clusters identified by single-cell consensus clustering (SC3) analysis are associated with cell types (colors) and enzymes used for reverse transcription (symbol types).

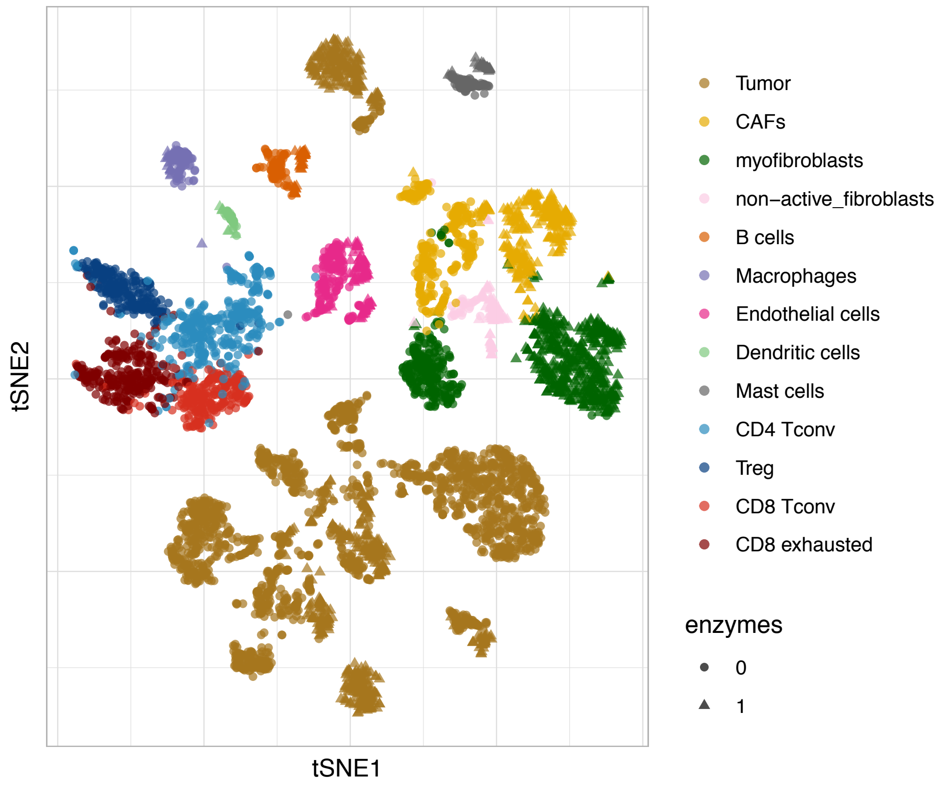


**Figure S13. Batch effect of enzyme treatment**
2D t-sne projection of all 5,712 single cells as in Figure 1A, with cells labeled by types (colors) and enzymes used for reverse transcription (symbol types).

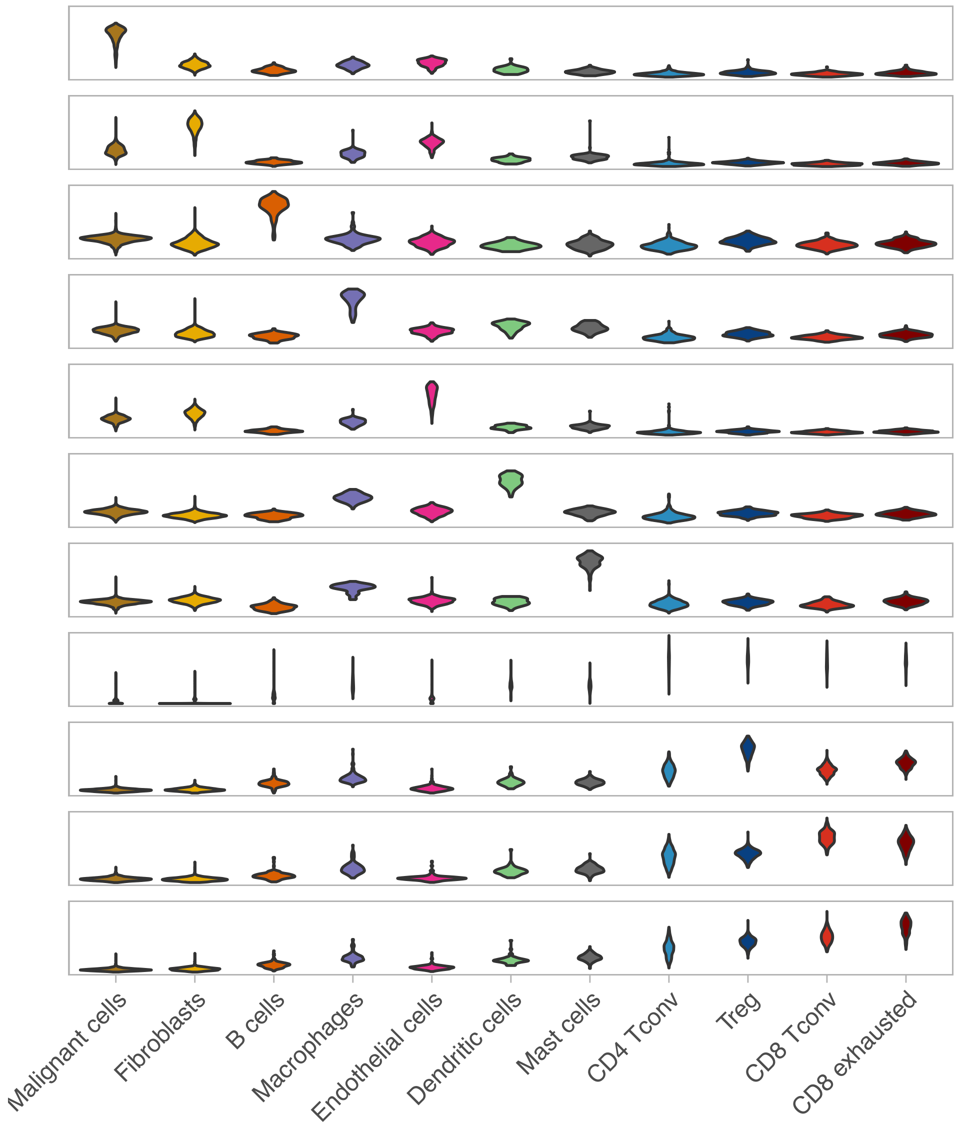


**Figure S14. Expression of DE markers (T1) across all cells stratified by cell types**

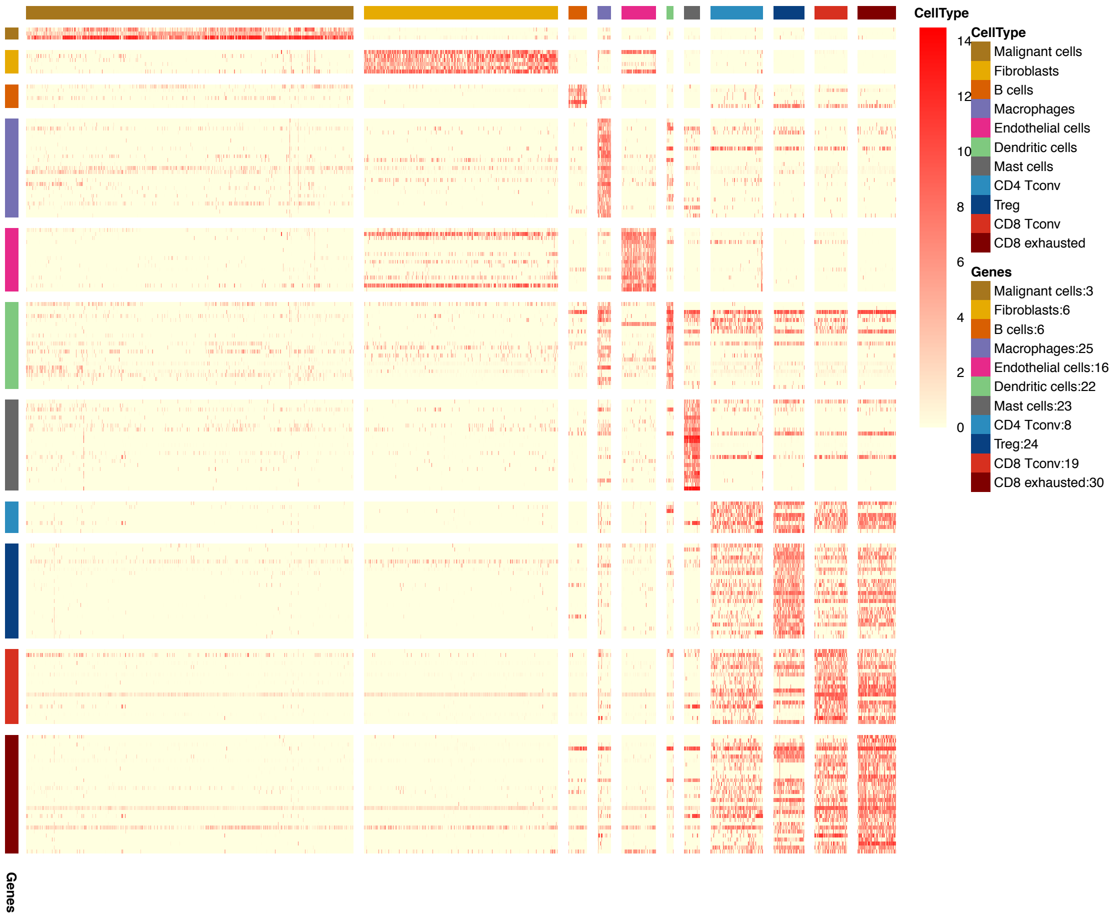


**Figure S15. Expression of genes shared between C2+T1 and LM22+C1 across all single cells stratified by cell types**
